# Model for breast cancer diversity and spatial heterogeneity

**DOI:** 10.1101/276725

**Authors:** J. Roberto Romero-Arias, Guillermo Ramírez-Santiago, Jorge X. Velasco-Hernández, Laurel Ohm, Maribel Hernández-Rosales

**Affiliations:** Conacyt. Instituto de Física y Matemáticas, Universidad Michoacana de San Nicolás de Hidalgo, Morelia, Michoacán, Mexico; Instituto de Matemáticas, Universidad Nacional Autónoma de México, Juriquilla Querétaro, Mexico; School of Mathematics, University of Minnesota, USA; Conacyt. Instituto de Matemáticas, Universidad Nacional Autónoma de México, Juriquilla Querétaro, Mexico

**Keywords:** Cancer mutations,Yule-Furry processes, Gene diversity, Gene heterogeneity.

## Abstract

We present and analyze a growth model of an avascular tumor that considers the basic biological principles of proliferation, motility, death and genetic mutations of the cell. From a regulatory network analysis and an analysis of genomic data we identify two sets of genes-a set of six genes and a set of sixteen genes-that are believed to play an important role in the evolution of breast cancer. Considering that cancer cells shape the tissue microenvironment and niches to their competitive advantage, the model assumes that cancer and normal cells compete for essential nutrients and that the rate of the “driver” mutations depends on nutrient availability. To this end, we propose a coupling between the transport of nutrients and gene mutations dynamics. Gene mutation dynamics are modeled as a Yule-Furry Markovian process, while transport of nutrients is described with a system of reaction-diffusion equations. For each representative tumor we calculate its diversity, represented by the Shannon index, and its spatial heterogeneity, measured by its fractal dimension. These quantities are important in the clinical diagnosis of tumor malignancy. A tumor malignancy diagram, obtained by plotting diversity versus fractal dimension, is calculated for different values of a parameter *β*, which is related to the occurrence of driver mutations. It is found that when *β <* 1, tumors show greater diversity and more spatial heterogeneity as compared with *β >* 1. More importantly, it is found that the results and conclusions are similar when we use the six-gene set versus sixteen-gene set.

## I. INTRODUCTION

Nowadays there is no consensus over how cancer is initiated; however, it is known that tumor growth occurs in several stages. The accepted general view is that a cell must undergo several gene mutations before it becomes cancerous. There are two kinds of mutations: “passenger” and “driver”. The former are gene changes that do not affect cell fitness or contribute to cancer development, and they may appear and eventually vanish during any stage of tissue development. The latter are gene changes that are causally involved in cancer development, typically conferring a functional change as well as a somatic evolutionary advantage, and are believed to play a crucial role in cancer progression [1, 2]. Because of this, cancer development is the result of the gradual accumulation of driver mutations that enhance cell proliferation rate and inhibit cell death rate leading to tumor progression [3–5]. The detailed factors that drive these mutations are unknown. Nonetheless, there is a general agreement that environment and heredity play important roles in cancer initiation. Tumor progression mainly involves two types of genes: (i) oncogenes and (ii) tumor suppressor genes [6–9]. Oncogenes encode proteins that control cell proliferation and apoptosis [10]. Oncogenes can be activated by structural alterations resulting from mutation or gene fusion [11], by juxtaposition to enhancer elements [12], or by amplification. Translocations and mutations can occur as initiating events [13] or during tumor progression, whereas amplification usually occurs during progression. Activation of oncogenes by chromosomal rearrangements, mutations, and gene amplification confers a growth advantage or increased survival of cells carrying such alterations. All three mechanisms cause an alteration in the oncogene structure, an increase or a deregulation of its expression [14]. On the other hand, tumor suppressor genes normally prevent unrestrained cellular growth and promote DNA repair as well as cell cycle checkpoint activation and maintain the activity of every cell. In most cancers the “bad mutations” of tumor suppressor genes reduce functions and make cells grow without control, and eventually accumulate to form a tumor [6, 8]. In normal cells, however, hundreds of genes intricately control the processes of division and death, so that, growth is the result of a balance between the activity of those genes that promote cell proliferation and those that suppress it. Cancer cells originate within tissue and no longer respond to many of the signals that control cellular growth and death. As they proliferate they diverge ever further from normality. Over time, these cells become increasingly resistant to the molecular controls that maintain normal cells, and as a result, they divide more rapidly than their progenitors and become less dependent on signals from other cells. Cancer cells even evade programmed cell death, despite the fact that their multiple abnormalities would normally make them prime targets for apoptosis.

Phenotypic and functional heterogeneity usually arise among cancer cells within a tumor as a consequence of genetic variations, environmental differences, and irreversible changes in cellular properties. Cancer cell heterogeneity displays striking morphological, genetic, and proteomic variability and represents a great challenge to diagnosis, treatment, and drug resistance. Spatial variations in cell genetic profiles lead to altered microenvironments. These positional variations are visible through analysis of tissue pathology images [15]. There are unlimited numbers of genetic and epigenetic alternatives along with all types of environmental stress that contribute to tumor evolution. Because of this, it is extremely difficult the identification of a universal molecular mechanism at the center of cancer initiation and development. However, recent studies have demonstrated that heterogeneity is observed to varying extent across a wide variety of cancers, with the identification of both clonal and sub-clonal driver mutations [16–18]. A complex interplay of gene expression, DNA alterations, gene mutations and environmental conditions are believed to be the main factors that drive tumor heterogeneity [19]. The term *“spatial heterogeneity”* will be interpreted here as the accumulation of gene mutations in different cells clusters that are spatially distributed nearby the tumor periphery.

In this paper we analyze a quantitative model of growth of an avascular tumor that considers the basic biological principles of cell proliferation, motility, death and genetic mutations. From genomic data, we identify two sets of genes-a set of six genes and a set of sixteen genesthat are believed to play an important role in breast cancer tumor growth. On the other hand, it has been found that cancer cells shape the tissue microenvironment and niches to their competitive advantage [17, 20]. In fact, nutrients play the role of catalysts during the expression of genes leading to fluctuations and asymmetries in the gene propensities, [21–23]. Thus, our model incorporates nutrients as an environmental factor that catalyzes gene mutation dynamics. The transport of nutrients is described by a set of reaction-diffusion equations coupled to the stochastic gene mutation dynamics of each cell, modeled as a Yule-Furry Markovian process. Taking into consideration that quantitative measurements of diversity and spatial heterogeneity are important clues for clinical diagnosis of tumor malignancy, we calculate and analyze these properties by means of the Shannon diversity index and the fractal dimension. With these quantities we establish a tumor malignant-benign diagram for different values of parameter *β* that tunes the accumulation rate of driver mutations. It is found that for *β <* 1 the tumors display high genetic diversity with a Shannon index of 𝓗 *>* 3.5 and are spatially heterogeneous, whereas for *β >* 1 tumors develop less genetic diversity characterized by a Shannon index 𝓗 ≤3.5, and are spatially less heterogeneous. The paper’s layout is as follows: in section II we present the model with the set of reaction diffusion equations for the transport of nutrients as well as the equations for stochastic mutation dynamics. In section III we briefly explain the algorithms used to simulate the genes stochastic dynamics and the integration of the system of reaction-diffusion equations. In section IV the results of the numerical simulations of tumors and the analysis of the diversity and spatial heterogeneity is presented. Finally, in section V we discuss the results of the structure, diversity and heterogeneity of tumors as well as the possible applications of the present quantitative modeling as a means of cancer diagnosis.

## II. MODEL

Let us start with a reaction-diffusion model for the growth of an avascular tumor proposed by Ferreira *et al.* [24]. Tissue is made of three types of cells, namely: normal, cancer, and tumor necrotic cells, that live on a square lattice. Processes of proliferation, death and competition for nutrients among the normal and cancer cells are considered. Normal and necrotic cells may occupy one lattice site, however, more than one cancer cell can pile up at a given lattice site. Because of this, three field variables 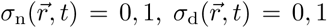 and 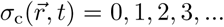are defined at each lattice site 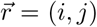 with 0<i, *j<L*, integer numbers. An initial cancer cell is placed at about the middle of the lattice. A nutrient supply (capillary vessel) is located horizontally at the upper side of the lattice. Periodic boundary conditions along the horizontal axis are defined. It is assumed that essential and nonessential nutrients diffuse from the capillary vessel towards each cell throughout the tissue (essential nutrients are glucose, amino acids, vitamins and minerals, and non-essential nutrients are oxygen, cholesterol and vitamins that are made naturally in the body). These nutrients are critical for DNA synthesis and for cell proliferation; therefore, they are considered important for the development of gene diversity [21–23]. Accordingly, we assume that high nutrient consumption leads to high driver mutation rates, while low nutrient consumption leads to a cell latent state with low driver mutation rates. This hypothesis, introduced as a stochastic term coupled to the reaction-diffusion Eqns (1-2), led us to show that the accumulation of driver mutations during tumor growth eventually yields high genetic diversity and spatial heterogeneity of tumor cells, the hallmarks of most cancers.

The reaction-diffusion equations that describe the transport of essential and nonessential nutrients for cell proliferation are the following [24]:

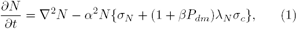

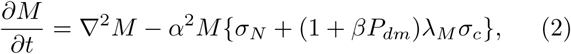

where 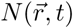and 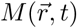are the concentrations of essential and nonessential nutrients, respectively. The parameter *α* represents the nutrient consumption rate for normal cells. The ability of cancer cells- enumerated by *σ*_*c*_-to outcompete normal cells (*σ*_*n*_) for essential and nonessential nutrients is denoted by *λ*_*N*_and *λ*_*M*_, respectively. The transport of nutrients on the right side of Eqns (1-2) include a term that is the product of two quantities, the parameter *β* and the probability *P*_*dm*_. The former describes the accumulation rate of driver mutations and the latter represents the probability of occurrence of driver mutations. Hence, the product *βP*_*dm*_quantifies the accumulation of driver mutations in each cancer cell. This stochastic term modifies the reaction term that accounts for the ability of cancer cells to compete for nutrients. Note that when *β* = 0, Eqns (1-2) reduce to the nutrient transport equations introduced in [24].

These transport equations are complemented with the probability of cancer cell division driven either by random mutations or by nutrient consumption, as well as the probability of death. This probability can be expressed as [24]:

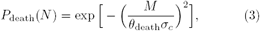

where *θ*_death_ controls the shape of this sigmoidal curve.

Cancer cell proliferation is modeled as a cell division process, *A*, which can occur in two independent ways, namely, (i) due to random driver mutations *A*_*M*_, or (ii) due to nutrients consumption, *A*_*N*_. Thus, the probability of cancer cell division can be written as the sum of two probabilities:

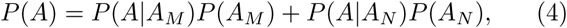

where *P* (*A |A*_*M*_) is the probability of division due to random driver mutations, and *P* (*A|A*_*N*_) is the probability of division catalyzed by nutrient consumption. By using Bayes’ property one can write, *P* (*A|A*_*M*_*)P* (*A*_*M*_) =*P* (*A*_*M*_*|A*)*P* (*A*) so that Eqn (4) can be recast as:

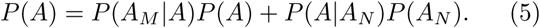

Solving for *P* (*A*) we obtain the probability of cancer cell division:

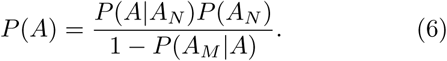

We now assume that the numerator of this equation catalyzes mutations by nutrient consumption as [24]:

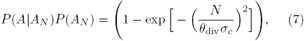

where *θ*_div_ controls the shape of this probability. By assuming that the probability of random driver mutations is *P* (*A*_*M*_*|A*) ≡ *βP*_dm_, the probability of cancer cell division, *P* (*A*) = *P*_div_, in Eqn (6) is given as

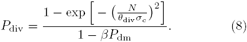

Since *P* div*≤* 1, then 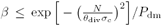; this means that driver mutations happen at a higher rate in those regions where nutrients concentration satisfies this inequality. Note that when the nutrient concentration is large the right side of this inequality is small whereas when nutrient concentration is small the right side of this inequality is large. Therefore, regions with high nutrient concentration favor cell survival and increase the driver mutation rate, nonetheless, regions with low nutrient concentration disfavor driver mutations. In this sense, nutrients play the role of a *“catalyst”* for driver mutations.

### A.Gene types

To determine which genes should be incorporated into the mutation dynamics, we analyzed data from the Data Release 23 from the European Union breast cancer project (BRCA-EU). We found a set of sixteen genes that are believed to play a major role in breast cancer development. These genes with their corresponding frequency of mutations are shown in Fig. 1. In this set we can identify the tumor suppressor genes TP53, ATR, ATM, E2F1; oncogenes: BRCA1, ERBB2, MDM2, NRAS, HRAF and kinase regulators CHEK2, KRAS, CHECK1, AKT1 and CDK2. On the other hand, a recent regulatory network analysis in breast cancer [25] inferred from gene expression data suggests that there are six genes, namely: TP53, ATM, ERBB2, BRCA1, MDM2 and CDK2, that play a major role in breast tumor development. This finding appears to be consistent with the general belief that successive driver mutations are produced mainly by six genetic variations [5, 6, 8, 9, 22, 25, 26]. Based on this data we focus our attention on two histograms: one with the sixteen genes –see Fig. 1– and the second with only the six genes from the regulatory network analysis [25], see inset Fig. 1.

**FIG. 1:**
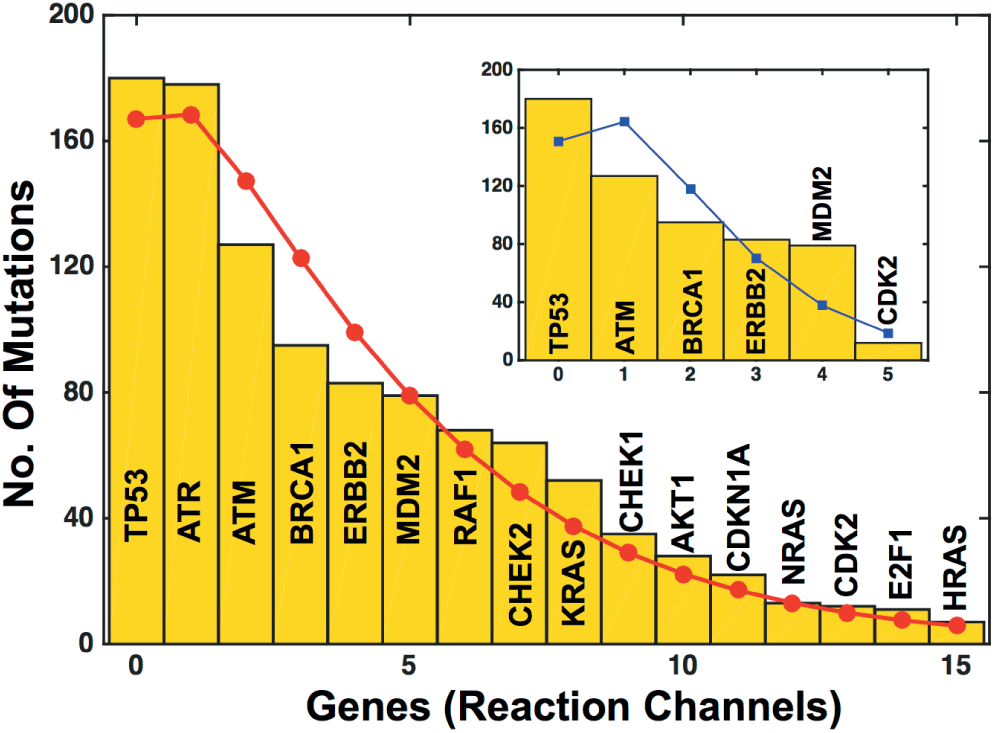
Histogram of the distribution of mutations of sixteengene set that are believed to play a major role in breast cancer development. Data is taken from Breast Cancer ICGC Project (https://dcc.icgc.org/) for the European Union. The inset shows the histogram of the distribution of mutations of six-gene set that play a crucial role in breast cancer as suggested from a regulatory network analysis [25]. The red line with circles and the blue line with squares represent the results of fitting a negative binomial distribution to each histogram. See text for more details.

Since mutation occur as a sequence of independent events (Markov chain), and because in each event a driver mutation may occur with probability *p* (success), accumulation of driver mutations can be described with a negative binomial distribution [27–29]. At this point it is important to note that a negative binomial distribution is commonly used to model gene expression in RNA sequence experiments [30–32]. Taking this into consideration, a negative binomial distribution with parameters, *p* (probability of success) and *r* (number of fails) is fitted to both histograms. The results of these fittings are shown with solid lines in Fig. 1. The sixteen-gene fitting shown in Fig. 1 with a red line and circles yielded the results *p* = 0.2578 ± 0.0136 and *r* = 1.3591 ± 0.0877 with a goodness of fit *Q* = 0.9301 and correlation coefficient *R* = 0.9779. Similarly, the result for the six-gene fitting is plotted in the inset of Fig. 1 with a blue line and squares. This fitting yielded the results *p* = 0.6560 ± 0.0481 and *r* = 3.1189 ± 0.6529 with a goodness of fit *Q* = 0.8713 and correlation coefficient *R* = 0.8699. This analysis indicates that the histogram made with either the set of gene mutations represents reasonably well the genomic data. The quantitative results of this fitting analysis will be used in the implementation of the stochastic simulations of the tumor growth dynamics (Section III A). To figure out if there are significant changes in the results when using either set of genes we carried out simulations using both sets. It is found that the results obtained with both sets are consistent and robust. Thus, in principle, it is sufficient to consider the six-gene set. For this reason most of our analysis is carried out with the six-gene set. In section IV we elaborate more about these findings.

### B.Mutation dynamics

By assuming that a driver mutation depends only on the previous cell genetic state, one can model the mutations dynamics by means of a Yule-Furry Markovian process. [33]. The corresponding master equation is

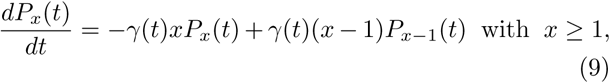

where *P*_*x*_(*t*) represents the probability that a given cell in the tissue undergoes *x* (*x* = 0, 1, 2,*…*) mutations at time *t*, and *γ*(*t*) *>* 0 is the hopping probability that one new mutation, *x → x* + 1, will happen in the time interval [*t, t* + *dt*). The solution of Eqn. (9) is a geometric probability distribution with argument,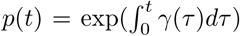 [33]. At this point we would like to relate the hopping probability to the microenvironment in tumor progression. To this end we consider recent findings in cancer development that suggest that interactions between cancer cells and their tissue habitat are reciprocal, and often times, cancer cells shape the tissue microenvironment and niches to their competitive advantage. From this point of view the tissue microenvironment is regarded as a complex system that leads to dynamic states with multiple components that influence cancer clone evolution [17, 20]. One of the simplest ways of incorporating this view is by assuming that nutrient spatial gradients modify in some way the acquisition rate of new driver mutations [34]. Thus, one can assume that there is a relationship between the concentration of essential nutrients and the hopping probability of acquiring new mutations, represented by γ(*t*). Taking this into consideration we make the *anzatz*, 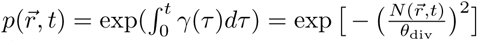 where 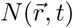 represents the concentration of essential nutrients at position 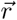 at time *t*, and *θ*_div_ is an adjusting parameter that controls the shape of the sigmoidal curve. Therefore, there is an intrinsic nonlinear coupling between the master equation that describes the mutation dynamics of each cell located at position 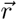 and the reaction-diffusion system that describes the nutrients concentration at any position in the tissue. Thus, to fully describe the dynamics of this complex system of equations we need to perform stochastic simulations (as described in section III).

It is known that driver mutations can be randomly activated by structural alterations resulting from mutation or gene fusion, by juxtaposition to enhancer elements, or by amplification of random mutations acquisition. These random activations can be modeled by a Poisson process distribution [6, 21, 35–37]. Then, one can write the total probability distribution of having a driver mutation, at a given cell in the tissue, at time *t*, as the product of two independent probability distributions, namely,

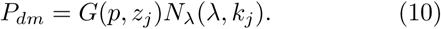

Here, *G*(*p, z*_*j*_) is a geometric probability distribution with mean (1*-p*)*/p*, where *p* is given by the *anzatz* indicated above, and *z*_*j*_ is the number of viable driver mutations of gene *j*. The function *N*_*λ*_ is a Poisson probability distribution with mean *λ* and represents the probability of occurrence of *k*_*j*_ driver mutations of gene *j* at a given cell. This factorization plays an important role in the implementation of the stochastic simulations to describe the tumor gene dynamics.

Now that we have defined the mutation dynamics probability distributions, let us estimate the upper bound for the parameter *β* that appears in Eqns (1-2). Recall that *β < p/P*_dm_, and that the mean value of Eqn (10) is (1 - *p*)*λ/p*. Then, one can obtain the following upper bound for *β*:

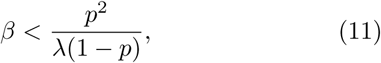

Since, *O*(1^*-*5^) *< p < O*(10^*-*1^) and *λ ∼ O*(10), *β* should be in the interval 0 *< β <* 10. These bounds for *β* will guide us to explore the following two limiting regimes:one in which the mutation rate of driver genes is low, when the cell is in a *“latent state”* because of low nutrients availability, and (ii) the other when there is a high rate of driver mutations. Because of this, one can think of *β* as a *“mutation driving parameter”.*

## III. NUMERICAL SIMULATIONS

Let us begin with a brief explanation of the algorithm that was used to simulate the genes stochastic dynamics.

### A.Stochastic Simulations

To simulate the stochastic mutation dynamics we used the Tau-Leaping Gillespie algorithm [38] which has demonstrated its usefulness in the simulations of different processes in molecular biology. In this approach, each gene can be thought of as a *reaction channel* in which the occurrence of a driver mutation is related to the reaction product –addition of a molecule– as a result of a chemical reaction. We consider each gene as a monomolecular reactant, so that the number of reaction channels equals the number of genes involved in the tumor evolution. Let us assume that at a given time *t*, the state of the system is defined by the vector **x**(*t*) = (*x*_1_(*t*),*…, x*_*n*_(*t*)), in which each coordinate represents the number of mutations (reactions) in each of the *n* genes. Then, the change of this state vector in the time interval [*t, t* + *τ)* is given as

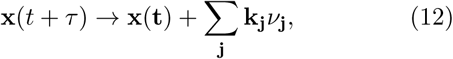

where **k_j_** is a vector of random numbers generated from a Poisson distribution with mean **a_j_**(**x**)*τ* and *ν***_j_** is the vector that increases the population at each channel, *j*, by 0, or 1. The channel selection, *j*, is made according to the negative binomial probability distribution obtained from the fits to the genomic data for either the set of sixteen or six genes whose results are presented in Fig. 1. In the simulations a few configurations are obtained in which the channel number is larger than the number of genes. In these cases the first gene type is chosen to be consistent with the fact that the first gene in the set is the one with a larger number of driver mutations. See Fig. 1.

Let **a_j_**(**x**) be the propensity functions that represent the probability of having one mutation at time *t* at any of the available channels. Since mutations occur with equal probability regardless of the gene (channel) then the values of **a_j_**(**x**) are equal to one for every channel. Therefore, the time *τ* required for the population of mutations to change in one unit is written as:

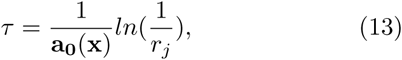

where 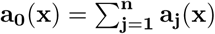 and *r*_*j*_ is a random numberuniformly distributed in the interval [0, 1] corresponding to gene *j*. Since the dynamics of the tumor evolution are the result of the coupling of the genes mutations and the nutrient dynamics, the extended version of the tau-leaping method is applied to obtain an effective sampling of the relevant quantities [39–42]. Thus, the change of the system’s state **x**(**t**) during a time *τ* occurs in accordance with the following equation

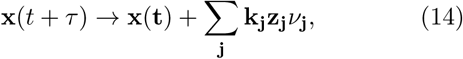

where **z_j_** is a vector formed with random numbers which are distributed according to a geometric distribution, and **k_j_** is the random vector generated from a Poisson distribution. We chose these random numbers distributions because driver mutations dynamics is described by the product of these two probability distributions; see Eqn (10).

### B.Numerical integration

The solutions of the reaction-diffusion system, Eqns (1-2), together with the probabilities for division and death, Eqn (8) and Eqn (3), respectively, are calculated numerically. We assume that normal, cancer and necrotic cells live on the sites of a square lattice of size *L×L* =500*×*500 [24]. Essential and nonessential nutrients for cell proliferation are continuously supplied through a capillary located at the top of the lattice simulating the bloodstream. Eqns (1-2) are integrated using zero flow boundary conditions at the left, right, and lower sides of the square domain. In order to obtain a homogeneous diffusion of nutrients, the equations are solved locally for each node populated with cancer cells, using a grid of size 10 × 10 units with zero flow boundary conditions, until the steady state was reached. After locally solving the equations, the reaction-diffusion equations are solved globally until a simulation cycle is completed. A simulation cycle consists of a complete sweep of the lattice, that is, once each site of the lattice has been visited. From now on one simulation cycle will be considered as a generation time, *T.* At the beginning of the simulations all cells are normal except for one that has developed cancer and is located near the lattice center. It is assumed that the initial cancer cell has suffered mutations in the gene TP53, which is the gene that plays a crucial role in tumor growth. In the simulations, cancer cells are chosen with the same probability. Once this initial cell begins to proliferate its descendants undergo mutations in all the other genes according to the probability distribution given in Eq (7). After a cell division occurs the daughter cell position is chosen randomly as one of the four nearest neighbors of the mother cell position. To this end the normal cell that was located at that site is removed. Then, we choose a random number *r* distributed uniformly in the interval [0, 1] and compare it with the probability *P* (*A|A*_*N*_*)P* (*A*_*N*_), that a mutation occurred (this probability is written in Eq (7)). A mutation occurs if *r > P* (*A|A*_*N*_*)P* (*A*_*N*_), otherwise it is rejected. A new cycle is initiated by randomly choosing a new cell and repeating the procedure for the mutation dynamics. To estimate the average time of a typical simulation, measured as the total number of cycles in a run, we carried out simulations of various lengths such that the mutation frequency of each gene was about its mutation frequency in the genomic data. See Fig. 1. This estimation led us to conclude that on average, a simulation of 800 cycles is sufficient for each gene to reach its mutation frequency of the genomic data. A typical simulation of this length yielded tumors with size smaller than 450*×*450 lattice sites for most combinations of values of the model parameters considered here. To understand the statistical meaning of the results we performed averages over 5, 10 and 20 simulations. It was found that the results were consistent within one standard deviation with those corresponding to just one simulation. Therefore, the results we report here correspond to one simulation of the system. The simulation parameters values were chosen in accordance with reference [24], that is, *θ*_div_ = 0.3, *θ*_death_ = 0.01, *λ*_*M*_ = 10, *λ*_*N*_ = *{*50, 100*}* and *α* = 4 × 10^*-*3^, 6 × 10^*-*3^. Since the values of the parameter *β* correspond to the mutation rates of driver genes we carried out simulations for *β* = 0.0, 0.25, 0.5, 0.75, 1.0, 2.0 and 4.0.

## IV. RESULTS

This section presents and discusses the results obtained from the numerical simulations of the model explained in section II. The dynamics of gene mutations are described in terms of the competition for nutrients between normal and cancer cells as well as the random dynamics of gene mutations that drive tumor progression. A thorough genomic data analysis yielded sixteen genes that are believed to play an important role in breast cancer development –see Fig. 1. In addition, we considered the results of a recent gene regulatory network analysis that took into account genetic and environmental aspects of breast cancer [25]. It was found that genes HER2(ERBB2), MDM2, TP53, as well as the regulatory genes HER2*→*TP53, *→*CDK2*→*BRCA1, ATM*→*MDM2, TP53*→*ATM are critical for breast cancer development. Based on this genomic data analysis, detailed simulations of tumor growth were carried out incorporating gene mutations dynamics with both sets of genes. Let us begin by considering the set of six genes obtained from the regulatory network analysis. Fig. 2 presents the tumor spatial distribution of mutations after one simulation cycle for each one of the six genes. The results have been arranged in a clockwise direction according to a decreasing number of mutations starting from the TP53 gene, which accumulates the most mutations. The central figure represents the superposition of the spatial distribution of mutations in the six genes under consideration in the tumor. An interesting feature to note in this result is the accumulation mutations at the upper tumor periphery. This is expected since the nutrient capillary supply is located at the top of the tissue domain, and the nutrient concentration gradient drives both cell division and mutation. The results indicate that genes TP53 and ATM undergo a major spreading, suggesting that cancer progression may be mainly due to the accumulation of mutations of these genes. The central part of the tumor (white region) represents the spreading of genes BRCA1 and ERBB2, which accumulate fewer mutations, suggesting that their contribution to the tumor progression is not important. Genes CDK2 and MDM2 accumulate mutations in a smaller region of the tumor close to its periphery; however, most of the tumor inner space (white region) shows no indication at all that these genes contribute to tumor growth. The observed spatial distribution of mutations in these figures suggests that tumor structure develops a certain degree of mutation diversity and spatial heterogeneity and it is similar to what is observed in the clinical analysis of biopsies [43–46]. The diversity and spatial heterogeneity will be quantified below. The tumors’ heterogeneity and diversity are quantified by using the k-means clustering algorithm [47] and Shannon entropy index [48], respectively. The k-means clustering algorithm yields a graphical distribution of cell clusters according to the number of mutations and is referred to the initial position of the tumor center of mass. To this end the following quantity is calculated [49],

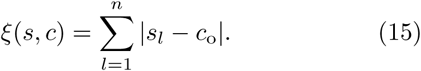

where *n* is the number of mutations in the *l*-th-cluster weighted with respect to the central cluster around the tumor center of mass, *c*_o_, and *s*_*l*_ is the distance of the *l*-th-cluster to center of mass. The Shannon entropy index is defined as 𝓗 = ∑_*i*_ *P*_*i*_*lnP*_*i*_ where *P*_*i*_ is the probability that *i* mutations occurred in the whole cancer tissue. To compute *P*_*i*_ we counted the number of cells that underwent one mutation, two mutations, three mutations, etc., and then, we divided this quantity by the total number of cancer cells. Therefore, the Shannon index is a measure of the diversity of cell mutations related to the number of mutations each tumor cell underwent. In ecology, Shannon index values lying in the range 1.5 *<𝓗<* 3.5 are considered as a normal diversity of species [48]. However, *𝓗>* 4 indicates a very rich community. In the present case an increase in the index of diversity, *𝓗*, is directly related to the abundance of gene mutations. It occurs either when there is a large number of cells with a relatively small amount of mutations or a few cells with a large amount of mutations. Clinically, the Shannon index can be measured through immunohistochemistry staining to evaluate cell-level heterogeneity as well as patients therapeutic response [50]. To quantitatively analyze the structure of cells clusters, the lattice sites (cells) were labeled according to their position in the vector formed by concatenating the columns of the square lattice. That is, the square lattice of size *L×L* is transformed into a one-dimensional lattice of length *L*^2^. Fig. 3 shows the cell clusters formation of a tumor obtained with model parameters values: *α* = 4*×*10^*-*3^, *λ*_*N*_= 100, and *β* = 1, after completing 800 cycles of simulation. The number of mutations of each cell is plotted against the position of each cell in the one-dimensional extended lattice. In Fig. 3(a) are shown the cells clusters and the corresponding tumor obtained with the six-gene set. The tumor has an index of diversity, *𝓗*= 5.15. Fig. 3(b) shows the cell clusters and tumor obtained with the sixteen-gene set; it is clear that it has a lower diversity index, = 4.93 as compared to tumor in Fig. 3(a). This is expected since as the number of genes increases the probability that each gene undergoes a number of mutations decreases because there are more genes available in which mutations may happen. Observe that the cell clusters that underwent the larger number of mutations are located at the tumor periphery, far from the tumor center of mass denoted by the symbol*×*. The spatially heterogeneous structure and high genetic diversity of the tumor is due to the occurrence of mutations in different genes at different tumor positions. Since the values of the diversity indices of the tumors obtained with both, the sixteen and the six-gene sets are similar, then the results are robust. Thus, using either set of genes yields results that are representative of the tumor genetic evolution. In Fig. 4 are shown the clusters obtained with the k-means clustering algorithm for each one of the six-gene set after *T* = 800 simulation cycles. The corresponding value of the index of diversity is also indicated. Observe that because the number of mutations of each gene is different then, the cluster structure varies across the tumor. Note also that the cluster corresponding to the gene that suffered the largest number of mutations has the highest diversity index. In fact, as the number of mutations decreases so does the diversity index. These results suggest that both the tumor suppressor gene TP53 and the oncogene BRCA1 play an important role in the structure of the tumor growth since they have high diversity index values, 𝓗= 3.97 and 3.64, respectively. As a matter of fact, genes with diversity index values greater than 3.50 are considered important attractors in the gene regulatory network analysis [25]. Thus, the tumor genetic structure and diversity index obtained from the present model is consistent with the gene regulatory network analysis. In addition, the ATM gene, which is considered to play a central role in the signal transduction of the early stages of tumor progression, has a diversity index 𝓗= 3.50, a value that is at the boundary between normal and high diversity. The other two genes, ERBB2 and MDM2, known as oncogenes have a diversity value that can be considered normal. The tumor suppressor gene CDK2 has the lowest diversity index, *𝓗* = 1.50. Figure 5 presents the cluster evolution of the distribution of mutations as well as the tumor progression at three stages of growth, focusing only on the mutations of the gene TP53. Note that the cells that underwent the largest number of mutations are always related to clusters that are on the tumor periphery, while cells that suffered few mutations are related to clusters located around the tumor center of mass. Note also that as the tumor size increases the diversity index increases too, suggesting that either a larger number of cells underwent a relatively small amount of mutations, or a few cells suffered a high amount of mutations. The diversity index of the first and second cluster configurations indicate a tumor with normal genetic diversity, while the third configuration has a diversity index value greater than 3.5 indicating a high genetic diversity. The same analysis was performed for the other five genes and the conclusions were similar. Figure 6 illustrates the cluster structure, left and right columns, obtained with the k-means clustering algorithm for the sixteen genes set after a simulation time of *T* = 800 cycles. The model parameters values used for these results are: *α* = 4*×*10^*-*3^, and *λ*_*N*_= 100 for the four values of *β*: (a) *β* = 0.25,(b) *β* = 0.75, (c) *β* = 2.0, and (d) *β* = 4.0. The corresponding tumors together with the color bar as a reference are plotted at the middle of the figure. Note that as *β* increases the tumor diversity index decreases. These results suggest a close relationship between the cluster structures and the branched structure of the corresponding tumor. The tumor regions that suffered more mutations are positioned at the tumor periphery whereas the regions that underwent few mutations are located close to the tumor center of mass. Frame (a) shows the cluster structure for *β* = 0.25 which has a diversity index *𝓗* = 4.99, an indication of high genetic diversity. Observe that the largest number of mutations occurs right above the cluster’s center of mass, indicated by *×*. However, a large number of mutations also occur at both sides of the tumor center of mass. As a consequence, the tumor becomes branched with the top region (yellow part) indicating the piece of the tumor where cells underwent the highest number of mutations. The cluster shown in frame (b) corresponds to *β* = 0.75 and has a diversity index 𝓗 = 5.0. In this case the region with the largest number of mutations is positioned at the upper right side, again in the tumor periphery. The cluster shown in frame (c) corresponds to *β* = 2.0 and has a diversity index 𝓗 = 4.39. There one sees that the number of mutations decreases by about half compared to the number of mutations accumulated in the clusters in frames (a) and (b). Notice that right above the cluster’s center of mass there is a bifurcation of two clusters that represent two regions of the tumor that suffered a large number of mutations. In this case the tumor developed fewer branches and became more compact at the middle. Frame (d) shows the cluster obtained for *β* = 4.0, which has a diversity index value 𝓗 = 3.42. For this value of *β* the cluster became less branched and more compact and homogenous, indicating that most of the tumor cells suffered about the same low number of mutations. Because of this, the tumor diversity index is the smallest of the four, which suggests that most of the cells conserved their genetic linage during proliferation. Considering altogether the clusters shown in Figs. 3 and 6, one sees that the cluster that shows the highest diversity corresponds to *β* = 1. More importantly, from the analysis of Fig. 6, one finds that the larger the value of *β*, the smaller the number of mutations while the tumor structure becomes less branched and more compact. This means that most of the tumor cells preserved their genetic lineage. At this point it is important to recall that in the present model the role of the microenvironment in tumor development is accounted for by considering the competition for essential nutrients for cell proliferation between cancer and normal cells. In fact, the nutrients transport equations, Eqns (1-2), were written in terms of the parameters *α* and *λ*_*N*_that measure, respectively, the nutrient consumption rate of normal cells and an additional factor by which nutrient consumption by cancer cells differs from their normal counterparts. Usually *λ*_*N*_ is chosen to be greater than 1 so that cancer cells consume essential nutrients at a higher rate than normal cells [24]. Fig. 7(a)-(d) shows the cell clusters as well as the full tumor spatial distribution of mutations for four combinations of values of the parameters *α* and *λ*_*N*_. The clusters and tumors have been referred to a coordinate system whose vertical axis represents the values of *α* while the horizontal axis represents the values of *λ*_*N*_. The *mutation driving parameter β* has been assigned the value *β* = 1, since for this value the tumor develops high diversity. It was found that for *α* = 4*×*10^*-*3^ and *λ*_*N*_= 50, the simulated tumors are compact and the cells located at the tumor periphery undergo up to 500 mutations, as indicated by the cluster analysis, Fig. 7(a). However, the tumors become fingerlike for *α* = 4 × 10^*-*3^ and *λ*_*N*_= 100 –Fig. 7(b). Here, the number of mutations decreased and the tumor main core became smaller than in the previous case. The cluster structure indicates that at the periphery (top right) there is a small cluster which is related to the tumor top branch, indicating that between 200 and 400 mutations occurred. The diversity index values are larger than four which indicates high genetic diversity, and corresponds to the high number of mutations that tumor cells suffered. For *α* = 6 *×*10^*-*3^ and *λ*_*N*_= 50, 100 –Fig. 7(c) and (d), respectively, the tumor and cluster structures become compact and decrease in size. In these cases the number of mutations that occur is a fraction (between 0.2 and 0.3) of the accumulation of mutations of the clusters in Fig. 7(a)-(b) and the diversity index value become smaller than four, which indicates normal genetic diversity. Note that as the number of mutations decrease, the cluster and tumor became smaller and more compact so that the index 𝓗 decrease.

**FIG. 2:**
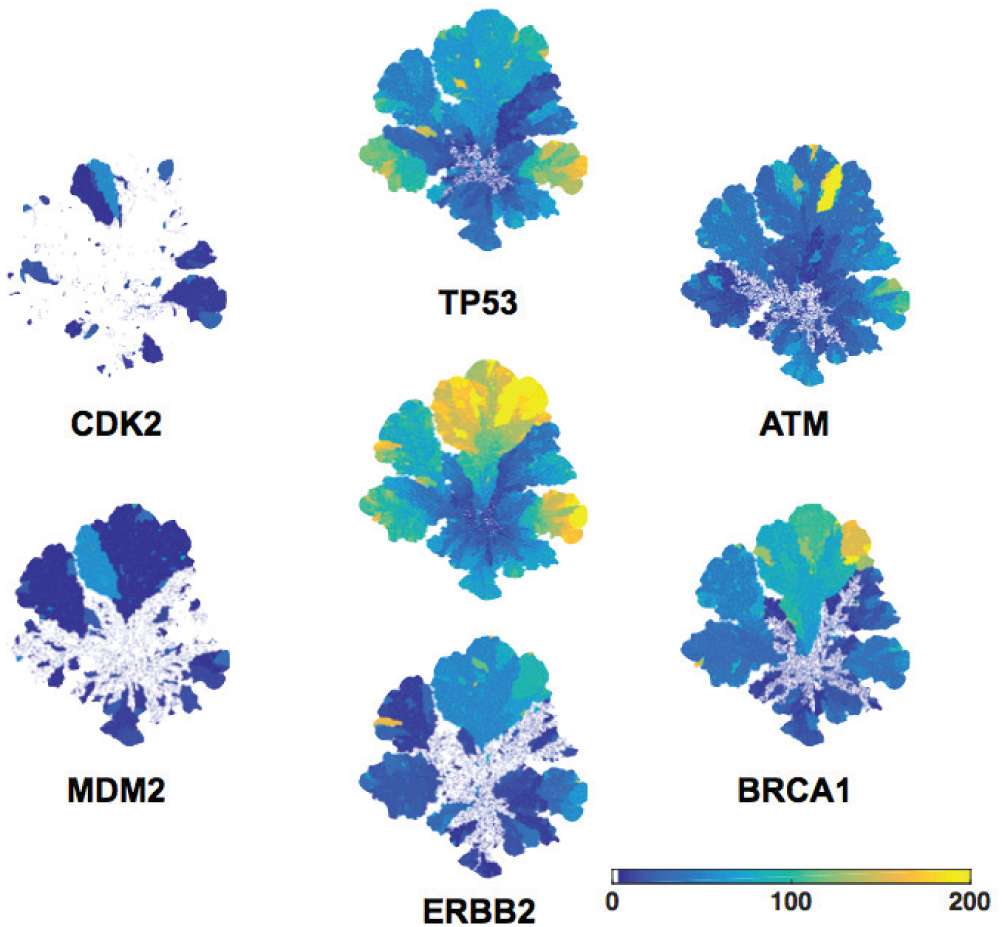
Starting from the top cluster and going in the clockwise direction are shown the spatial distribution of mutations. They are ordered in a decreasing way according to the number of mutation for each of the six genes (TP53, ATM, BRCA1, ERRB2, MDM2 and CDK2) indicated in the inset of Fig. 1. At the center lies the tumor showing the spatial distribution of mutations of the six genes altogether. The results correspond to the parameter values: *α* = 4*×*10^*-*3^, *λN* = 100, and *β* = 1, after *T* = 800 cycles. The scale of colors is related to the number of driver mutations for each gene.

**FIG. 3:**
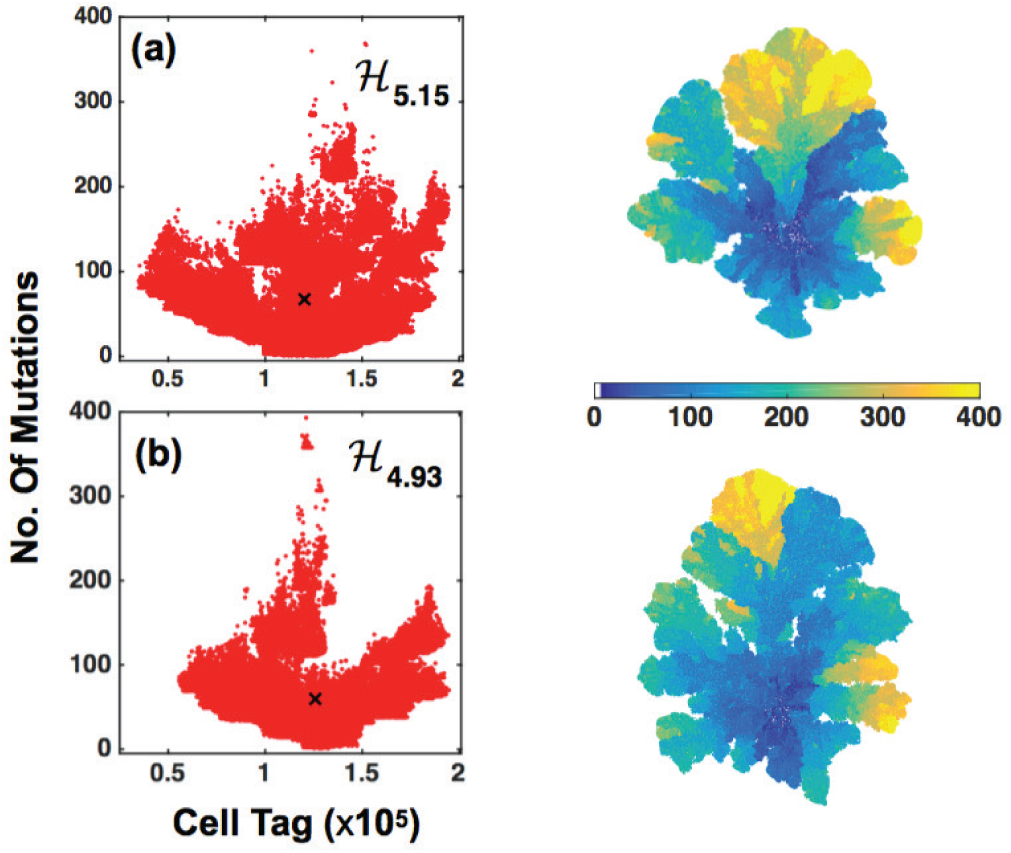
Clusters obtained with the k-means clustering algorithm for the same model parameters and time as in Fig. 2. (a) For the six genes set we obtained a diversity index, 𝓗 = 5.15. (b) For the sixteen genes set we obtained a diversity index, 𝓗 = 4.93. These values correspond to a high diversity tumor. Note that cells with a high number of mutations are located in the tumor periphery, far from the tumor center of mass indicated with the symbol *×*. The scale of colors is related to the number of driver mutations for each gene.

**FIG. 4:**
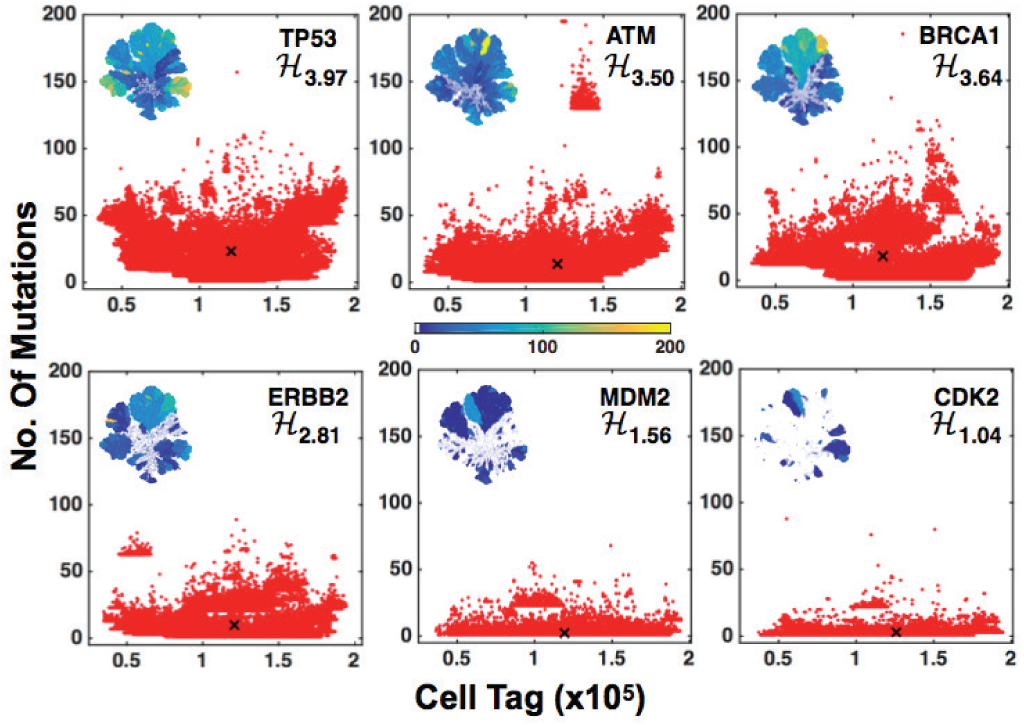
In red are the clusters obtained with the k-means clustering algorithm for each gene in the six-gene set. Note that the clusters structure that signals a large number of mutations are located far from the tumor center of mass (×). The inset in each figure represents the tumor with the corresponding spatial distribution of genes. Observe that the diversity index 𝓗 decreases as the cluster size decreases. These results were obtained for the same parameter values as in Fig. 2. The color bar at the middle of the centered column measures the number of mutations.

**FIG. 5:**
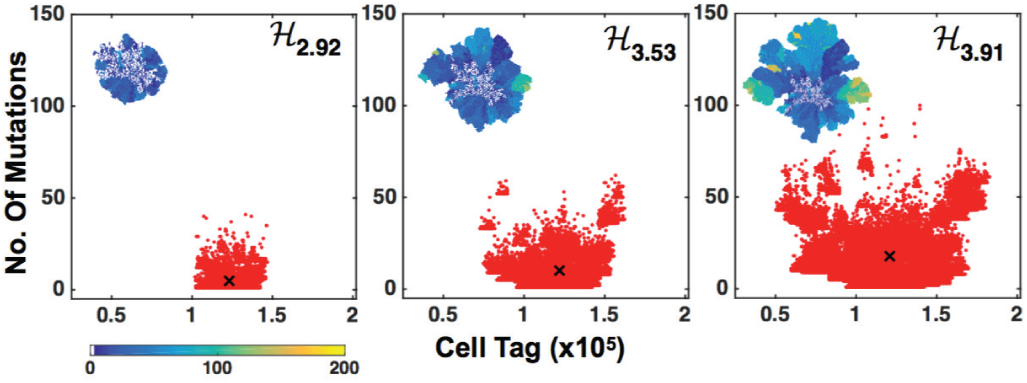
Cluster evolution showing the distribution of mutations of gene TP53 at three stages of the tumor growth: (a) *T* = 200 cycles, (b) *T* = 400 cycles, and (c) *T* = 600 cycles. On the upper left side of each frame are the spatial distributions of mutations of gene TP53 in the full tumor. The color bar indicates the number of mutations. Note that the diversity index increases from 2.92 to 3.91. These results were obtained for model parameters as in Fig. 2.

**FIG. 6:**
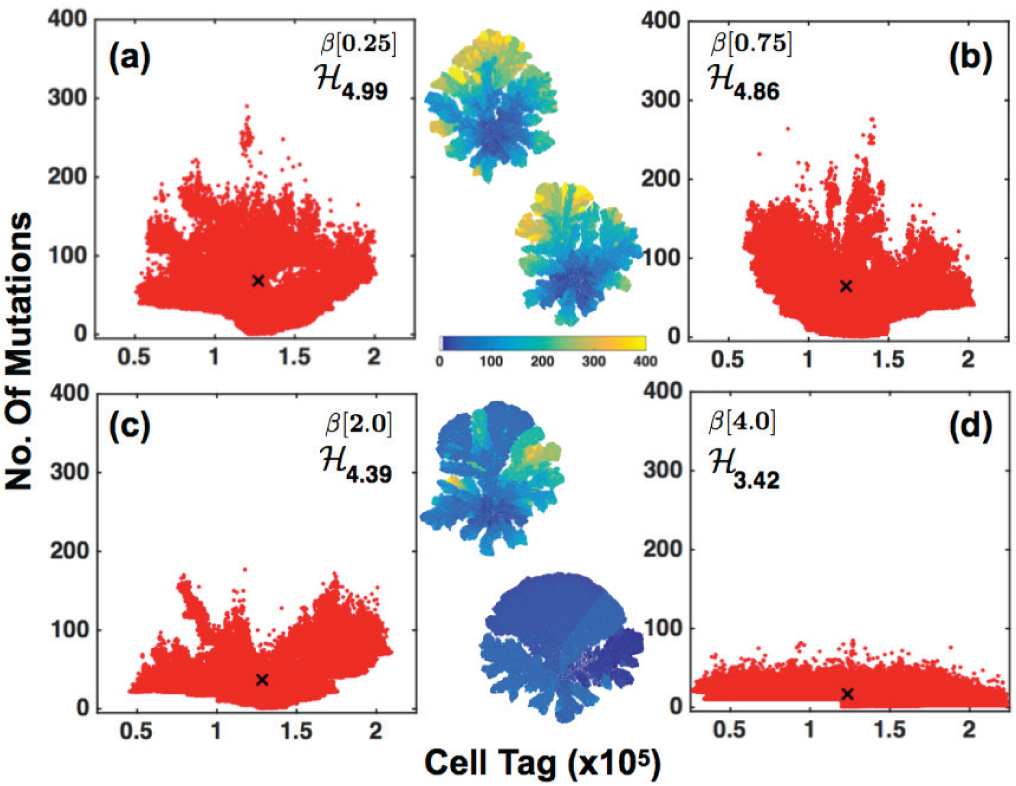
Tumor (at the middle column) and cluster structure (left and right columns) obtained with the k-means clustering algorithm after a simulation time *T* = 800 cycles for four values of *β*: (a) *β* = 0.25, (b) *β* = 0.75, (c) *β* = 2.0, and (d) *β* = 4.0. Notice that as the parameter *β* increases the diversity index 𝓗 decreases. These results are obtained for the model parameter values *α* = 4 10^*-*3^, and *λ* _*N*_ = 100. Note that tumor shows a branched structure with the top region (yellow part) indicating the cells located there underwent a large number of mutations.

**FIG. 7:**
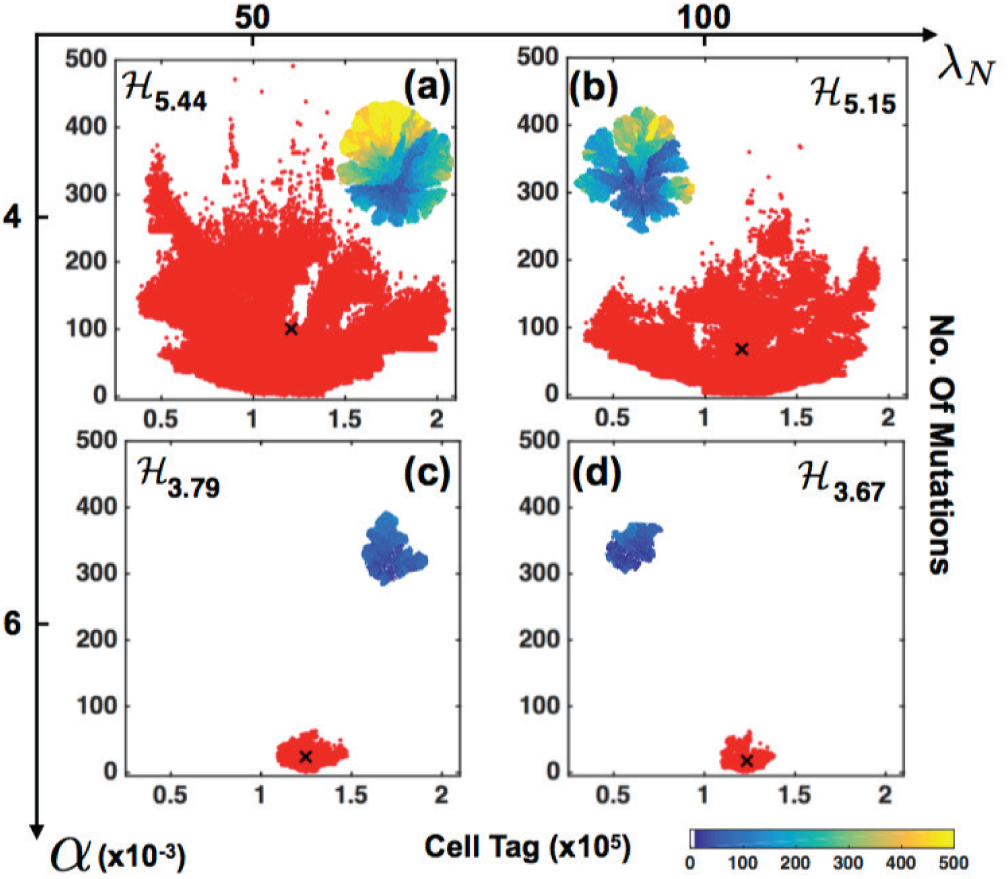
Cluster structure and spatial distribution of mutations in the tumor for four representative combinations of values of the parameters *α* and *λ*_*N*_, for *β* = 1, and simulation times *T* = 800 cycles. (a) *α* = 4 × 10^*-*3^ and *λ*_*N*_ = 50, (b) *α* = 4 × 10^*-*3^ and *λ*_*N*_ = 100, (c) *α* = 6 × 10^*-*3^ and *λ*_*N*_ = 50, and (d) *α* = 6 × 10^*-*3^ and *λ*_*N*_ = 100. The values of the index of diversity are also written for each case. Notice that these values decrease as the tumor becomes more compact and smaller in size. The color bar represents the number of mutations.

Another quantity that characterizes the tumor structure is the fractal dimension (FD). This quantity can be measured in histopathology slides of tissue samples and is an important step in the diagnosis of the malignancy of the tumor [51–53]. In addition, the change in texture or appearance of distortions in breast cancer tumors can be detected from mammograms by estimating the FD [54]. With the aim of relating these clinical measurements with the tumor structure and the spatial distribution of mutations of the in-silico tumors –for instance those presented in Fig. 2– the time evolution of the FD was calculated by means of the standard box-counting algorithm (Fig. 8). It was found that for each of the six genes, the FD increases monotonically as a function of time. The FD also becomes systematically greater for those genes that underwent more mutations, as expected. That is, tumors become more diverse and heterogeneous when one or more genes suffer many mutations in cells located at different positions. To express the FD time evolution in terms of a biological time scale one can relate one simulation cycle *T* with a biological cell division cycle that lasts about 35 hours [55]. Since a full simulation of the in-silico tumors lasts about 800 cycles, the typical simulations reported here correspond to approximately 28,000 hours, about 38.9 months of real time. The results shown in Fig. 8 indicate that the FD of the spatial distribution of the genes TP53, ATM and BRCA1 become asymptotically closer to each other for times longer than 12 months (*T >* 300 generations). In fact, the trend shown in the figure suggests that the FD behavior of the genes TP53, ATM, BRCA1, ERBB2 and MDM2 will asymptotically collapse onto one single curve for times *T >* 800 cycles, or longer 38.9 months of real time. More importantly, the FD evolution of the whole tumor, represented in the figure by a solid line, is quantitatively similar to the FD behavior of the spatial distribution of mutations of gene TP53, which is the gene that accumulates the most mutations. This result indicates that the fractal structure of the whole tumor is fully determined by the fractal structure of the spatial distribution of mutations of the gene TP53, which is thought to be the gene that plays a crucial role in cancer progression. The inset of Fig. 8 shows the time evolution of the whole tumor FD for five representative values of the *mutation driving parameter β*. Notice that for *β* ≤ 1, the FD follows approximately one single curve; however, for *β >* 1, the FD becomes systematically larger. This is not surprising since for smaller values of *β* more mutations occur in the tumor, and for larger values of *β*, the number of mutations in the tumor decreases. The results presented in the inset of Fig. 8 suggest that as the number of mutations increases, the time evolution of the tumor FD approaches a single universal curve.

**FIG. 8:**
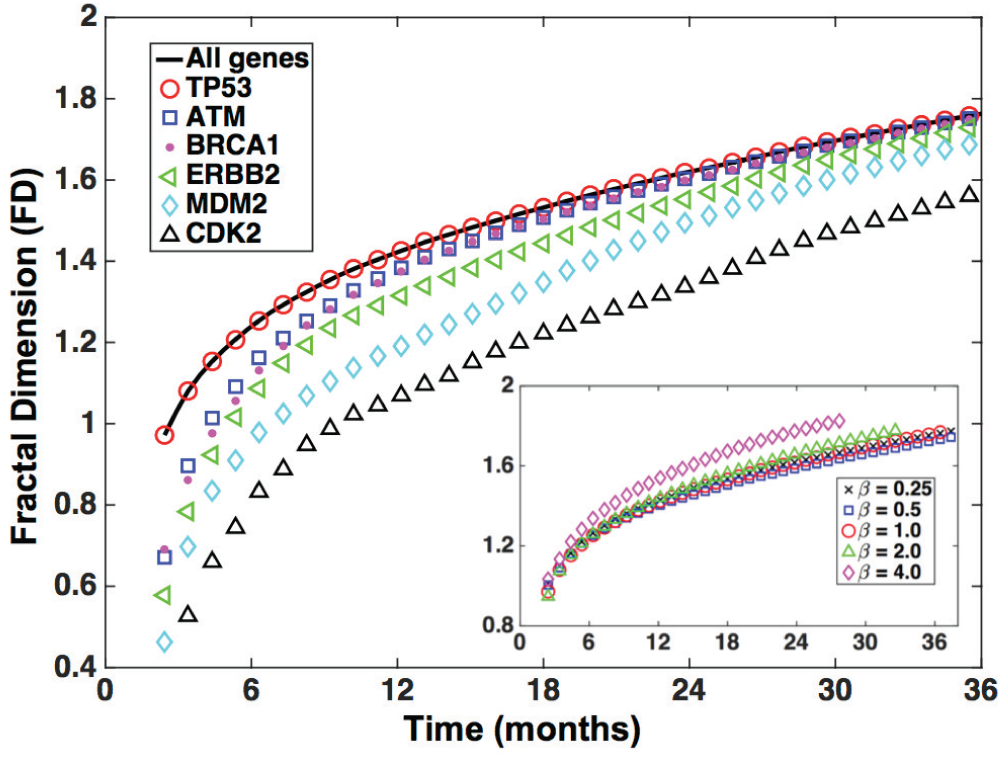
FD time evolution of the spatial distribution of each gene. Here we have considered the dynamics of the six-gene set. The symbols represent the time evolution of the FD corresponding to the spatial distribution of mutations of each gene. Note that the spatial distribution of mutations of gene TP53 has a FD time evolution that is quantitatively similar to that of the whole tumor, represented by the solid black line. These results correspond to a tumor with model parameters: *α* = 4 × 10^*-*3^, *λ*_*N*_ = 100 and *β* = 1. The inset shows the time evolution of the whole tumor FD for different values of *β* and same values of *α* and *λ*_*N.*_ These results indicate that as the number of mutations increase, the time evolution of the tumor FD approaches one universal curve.

In silico tumors have been generated for the model parameters *α* = 4 × 10^*-*3^, *λ*_*N*_= 100, different values of the *mutation driving parameter β*, and the six-gene mutation dynamics. For each of them, the diversity index, 𝓗, and the FD have been calculated. With the results of these simulations, a “2D malignancy diagram”, 𝓗 versus FD, has been calculated. The results are shown in Fig. 9. There one sees that 𝓗increases monotonically as a function of FD for all values of *β*. Note that with the exception of the curve corresponding to *β* = 0.5, all the other curves intersect at the crossing point of the dashed lines with coordinates *FD* = 1.3, *𝓗* = 3.5. This behavior suggests that there is a“critical point” in the tumor structure as a result of the spatial distribution of mutations. Recent studies have found that when FD *<* 1.3, breast cancer tumors are benign, while for FD *>* 1.3, tumors become malignant [54]. In addition, it is known that when the diversity index is in the range 1.5 *<𝓗<* 3.5 the tumor genetic diversity is considered normal, while for 𝓗 *>* 3.5, the tumors genetic diversity becomes high. Taking these clinical results into account, one can divide the plane versus FD into four quadrants with the axis crossing point located at *FD* = 1.3, *𝓗* = 3.5 as indicated in Fig. 9. The (*FD, 𝓗)* points located in the upper right quadrant *FD >* 1.3 and *>* 3.5 correspond to a malignant tumor. However, points located in the lower left quadrant (*FD <* 1.3 and 𝓗 *<* 3.5) correspond to a benign tumor. The inset of Fig. 9 shows the malignancy diagram obtained with a sixteen-gene mutation dynamics for three values of *β*. The diagram looks similar to that obtained with a six-gene dynamics, as shown in Fig. 9. All these results indicate that the predictions of our model are robust regardless of the use of six or sixteen-gene mutation dynamics. In addition, they are quantitatively consistent with clinical and experimental findings as indicated above.

**FIG. 9:**
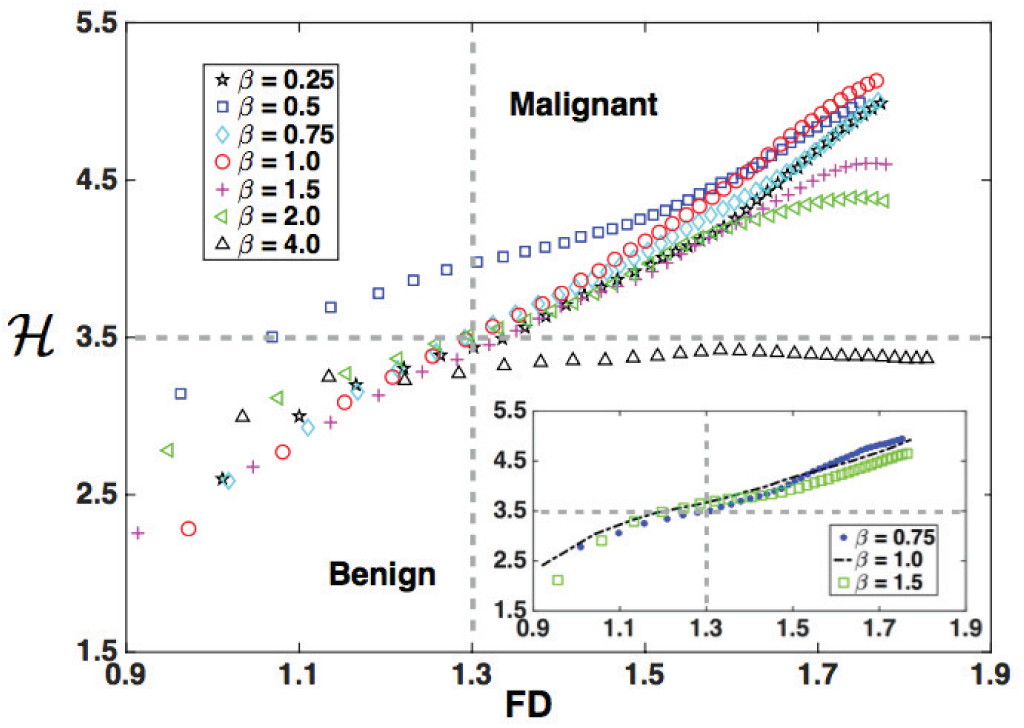
Diversity index 𝓗 versus FD for several values of *β*. The dynamics of the six genes set has been considered here. The results were obtained for a tumor with the model parameters: *α* = 4 × 10^*-*3^, *λ*_*N*_ = 100. Note that the curves intersect at (1.3, 3.5). Points in the upper-right quadrant correspond to a malignant tumor while those in the lower-left quadrant correspond to a benign tumor. The results shown in the inset correspond to the gene dynamics with sixteen-gene set and the results are consistent with those obtained with the set of six genes.

## V. CONCLUSIONS

We have presented and analyzed a quantitative growth model of an avascular tumor that considers the basic biological principles of cell proliferation, motility, death, transport of nutrients and gene mutation dynamics. We postulate that the gene mutation rate depends on both randomness and microenvironmental factors, such as essential and nonessential nutrient concentrations for cell proliferation. It was found that higher concentrations of nutrients is an advantage that favors cancer cell proliferation as well as a high accumulation of driver mutations, which in turn leads to genetic diversity and tumor heterogeneity. Gene mutation dynamics considered two sets of genes, one with six genes from a regulatory network analysis and the other with sixteen genes, from an analysis of genomic data, which are believed to play a crucial role in cancer progression. The coupling of mutation dynamics to microenvironmental factors was done by introducing a parameter, *β*, together with a probability distribution that regulates driver mutation dynamics. The mutations in turn defines the diversity and heterogeneity of the tumor. For *β <* 1, the rate of accumulation mutations is high and leads to high tumor gene diversity, whereas for *β >* 1, the rate of accumulations mutations is low and the tumor diversity becomes normal. For a given tumor one can calculate the diversity index,𝓗, and the FD for different values of the parameter *β*. Thus, a “malignancy diagram” based on 𝓗 versus FD was calculated Fig. 9. With the exception of the curve corresponding to *β* = 0.5, all the other curves intersect at the crossing point *FD* = 1.3, *𝓗*= 3.5 suggesting that this point indicates a critical change in the tumor behavior. More importantly, the results presented here suggest that the predictions of our model are robust whether we use six or sixteen-gene sets for the mutation dynamics. In addition, our findings suggest that tumor fractal structure and diversity are fully determined by the heterogeneous spatial distribution of mutations of gene TP53, which is though to play a crucial role in cancer progression. Finally, it is important to indicate that the predictions of our model can be quantitatively related to clinical and experimental observations.

## Authors contribution

JRRA, GRS and JXHV developed the model. JRRA, GRS carried out the simulations, the analysis of the results and wrote the paper. LO developed the preliminary version of the code for the tumor growth and proofread the final version of the manuscript. MHR provided the genomic data.

## Acknowledgments

JRRA would like to thank financial support from CONACyT-FORDECyT under grant number 265667. GRS would like to thank financial support from DGAPAUNAM grant number IN108916. JXVH would like to thank financial support from DGAPA-UNAM grant number IN110917. JRRA and GRS would like to thank enlightening discussions with Prof. Enrique HerńandezLemus. We would like to thank Carlos González-Castro for his technical support in the development of the stochastic simulations code. We would also like to thank Andres García-García for contributing to the genomic data search and sorting as well as a preliminary analysis.

